# Comparative RNA-seq based transcriptomic analysis of *Aspergillus nidulans* recombinant strains overproducing heterologous glycoside hydrolases

**DOI:** 10.1101/241273

**Authors:** Felipe Calzado, Mariane P. Zubieta, Gabriela F. Persinoti, Cesar Rafael Fanchini Terrasan, Marcelo V. Rubio, Fabiano J. Contesini, Fabio M. Squina, André Damasio

**Author notes:** These authors equally contributed to this work (Felipe Calzado, Mariane P. Zubieta). **To whom correspondence should be addressed:** André R. L. Damasio; Department of Biochemistry and Tissue Biology, Institute of Biology, University of Campinas (UNICAMP), Campinas-SP, Brazil.

## Abstract

Filamentous fungi are important cellular factories for the production and secretion of homologous and heterologous enzymes such as carbohydrate-active enzymes. However, the regulation of protein secretion in these microorganisms requires more profound studies since the enzyme levels produced are usually below the levels required by industry for profitable processes. Genomic and transcriptomic approaches have been used to understand the overexpression and production of heterologous enzymes and their capacity to induce different cellular biological processes. To investigate this regulation, *Aspergillus nidulans* recombinant strains were analyzed by transcriptomics. We designed three *A. nidulans* recombinant strains producing the following heterologous proteins: alpha-arabinofuranosidase (AbfA), beta-glucosidase (BglC) and thermophilic mannanase (Tp-Man5). The heterologous genes *abfA* and *bglC* were highly expressed, while *tp-man5* mRNA levels were low and similar to those of a reference gene. There was an indirect relationship between mRNA and protein secretion levels, suggesting that transcription is not a bottleneck for target gene expression in this system. Despite the distinct features of the recombinant proteins, 30 differentially expressed genes were common to all the recombinant strains, suggesting that these genes represent a general response to the expression of heterologous genes. We also showed that the early activation of the canonical unfolded protein response (UPR) pathway by *hacA* alternative splicing was normalized after 8h, except in the strain expressing BglC, suggesting either no accumulation of the BglC misfolded form or the presence of an alternative endoplasmic reticulum (ER) stress and UPR pathway. Finally, to focus our analysis on the secretion pathway, a set of 374 genes was further evaluated. Seventeen genes were common to all the recombinant strains, suggesting again that these genes represent a general response of *A. nidulans* cells to the overexpression of recombinant genes, even thermophilic genes. Additionally, we reported the possible genetic interactions of these 17 genes based on coexpression network calculations. Interestingly, protein improvements are nongeneric, and improvements in the production of one target protein are not necessarily transferable to another one. Thus, this study may provide genetic and cellular background and targets for genetic manipulation to improve protein secretion by *A. nidulans*.

## INTRODUCTION

Fungi belonging to the genus *Aspergillus* feature saprophytic lifestyles and have a high capacity to produce large quantities of extracellular enzymes from complex organic materials (Lowe and Howlett, 2012). This natural ability for efficient protein secretion has led to their biotechnological exploitation as cell factories for industrial enzyme production. Notably, a broad range of homologous and heterologous enzymes such as amylases, xylanases, and cellulases are produced by aspergilli, thus contributing to the expansion and growth of the enzyme market (Lubertozzi and Keasling, 2009). If this market continues to increase at an annual rate of 8.2% over a 5-year forecast period, it is expected to reach approximately $7.1 billion by 2018 (Reseach, 2014). *Aspergillus* is among the main enzyme-producing microorganism, and produces 30% of commercial enzymes, according to a list from the Association of Manufacturers and Formulators of Enzyme Products (AMFEP, 2015).

Despite the abovementioned advantages, there are still many challenges to overcome regarding the expression of target enzymes using fungal systems. The production of a particular protein requires high amounts of mRNA, efficient translation of the target gene, targeting of the protein to the secretion pathway (if secretion is desired), folding, posttranslational modifications, and low or no degradation of the target protein in the extracellular medium. Several strategies have been developed to optimize the quantity the quality of enzymes; these strategies include genetic engineering of promoters for high levels of mRNA (Hirasawa et al., 2018), preferential codon usage (Cripwell et al., 2017), removal of introns (Chesini et al., 2018), engineering signal peptides for enhanced protein secretion (Roongsawang et al., 2016), and the deletion of proteases that could degrade the products (Havlik et al., 2017). The wide variety of tools developed to achieve high yields of recombinant enzymes highlights how complex this pathway is in filamentous fungi. This complexity has resulted in a large body of work dedicated to this topic and the rapid development of new techniques to obtain enzymes. However, in recent decades, there have been no major published advances that boost the yields of biotech products produced by fungi (Nevalainen and Peterson, 2014; Meyer et al., 2016).

Misfolding and/or errors in the processing of recombinant protein in filamentous fungi is a critical bottleneck resulting in the elimination of these proteins by endoplasmic reticulum (ER) quality control (Guillemette et al., 2007; Pakula et al., 2016). Misfolded proteins alter cell homeostasis and proper ER function, resulting in ER stress. ER stress activates conserved signaling pathways such as the *unfolded protein response* (UPR) and *ER-associated protein degradation* (ERAD); these pathways upregulate genes responsible for restoring protein folding homeostasis in cells and degrading misfolded proteins in the cytosol by the ubiquitin-proteasome system, respectively (Heimel, 2014).

Manipulation of the UPR pathway and its components has been a common strategy to improve the production of heterologous proteins in filamentous fungi (Hayano et al., 1995; Valkonen et al., 2004; Xu et al., 2005). Many ER stress-induced UPR genes, including protein folding-related genes such as chaperones and foldases, have been coexpressed with a heterologous gene. However, the overexpression of chaperones usually does not increase the production of heterologous proteins. In *S. cerevisiae*, the overexpression of *bipA* increased the amount of extracellular prochymosin by over 20-fold, although the secretion of thaumatin was not significantly improved (Harmsen et al., 1996).

Genomic and transcriptomic approaches have been used to gain a deep understanding of the overexpression and production of heterologous enzymes and their capacity to induce UPR, providing valuable information on *Aspergillus* genes involved in the secretion and coordination between UPR and ERAD. The induction of UPR in *Aspergillus niger* was investigated by transcriptomic analysis to compare the UPR induced by chemicals and that induced by the overexpression of tissue plasminogen activator (t-PA) (Carvalho et al., 2012). Approximately 94 genes were commonly induced, most related to the functional categories of protein folding, translocation/signal peptidase complex, glycosylation, vesicle trafficking, and lipid metabolism (Carvalho et al., 2012). In addition, another study showed that UPR results in the activation of approximately 400 genes in *S. cerevisiae*(7-8% of the genome) (Travers et al., 2000). This reflects the complex network of interactions between UPR and other signaling pathways in the cell. Transcriptome profiles can be particularly important for providing an overview of all genes and pathways regulated by the UPR in a cell.

Interestingly, protein improvements are highly variable, and improvements in one protein are not necessarily transferable to others. In addition, decades of rational and nonrational strains improvements have not been available to academia due to commercial confidentiality (Meyer et al., 2016). Here, we performed a comparative transcriptome analysis of three *A. nidulans* recombinant strains producing the following heterologous proteins: 1) GH51 alpha-arabinofuranosidase (AbfA) (*A. nidulans_AbfA_*); 2) GH3 beta-glucosidase (BglC) (*A. nidulans_BglC_*) and 3) GH5 thermophilic mannanase (Tp-Man5) (*A. nidulans_Tp-Mans_*). These enzymes have very different domains and posttranslational modifications, so our interest was to identify common differentially expressed (DE) genes that potentially represent a general cellular adaptation to the overexpression of heterologous genes, including a hyperthermophilic gene. The transcriptional profiles of *A. nidulans* recombinant strains were determined at early and late periods of induction. Despite the distinctive features of the recombinant proteins, 30 DE genes were common to all the recombinant strains. It is likely that these genes represent a general response to the expression of heterologous genes. Furthermore, 17 genes specifically related to the secretion pathway were DE in the recombinant strains suggesting again that these genes represent a general response of *A. nidulans* cells to the overexpression of recombinant genes, even a thermophilic gene. Our results provide new potential targets for genetic manipulation and the improvement of protein secretion yields in this microbial system.

## RESULTS AND DISCUSSION

### Recombinant proteins are secreted at different levels

To study the production of heterologous proteins in *A. nidulans*, three recombinant proteins were monitored: AbfA and BglC were cloned from *Aspergillus fumigatus*, and thermophilic Tp-Man5 was cloned from the hyperthermophilic bacterium *Thermotoga petrophila*. These target sequences were cloned into the pEXPYR vector and transformed into *A. nidulans*_A773_(Segato et al., 2012). Interestingly, the recombinant proteins were secreted at different levels, *i.e*, AbfA > BglC > Tp-Man5. The transformed strains were denominated *A. nidulansAbfA, A. nidulans*_BglC_ and *A*. nidulansTp-Man5.

Initially, the time course of recombinant enzymes production was evaluated (Figure 1). Despite differences in the secretion of recombinant proteins, the profiles of mycelium dry weight, amount of extracellular proteins and final pH were very similar in the recombinant strains. The similarities among these parameters indicate that the overexpression of heterologous genes may not imply major physiological changes in the recombinant strains. AbfA showed the highest level of secretion, with an activity peak at 48 h (9.53 U/ml) (Figure 1A; *middle panel)*. BglC secretion was lower than AbfA and the activity peak was early at 24 h (Figure 1B; *middle panel*; 8.73 U/ml). Overall, after 48 h of cultivation, a protease degradation profile was observed for each of the recombinant strains (Figure 1; *bottom panel)*. At this cultivation time point, cells probably undergo disruption due to the absence of nutrients. Higher beta-glucosidase activity was detected from 72 to 120 h (Figure 1B; *middle panel*), corresponding to a native intracellular beta-glucosidase (AN2828) whose amino acid sequence/identity was further confirmed by LC-MS/MS (data not shown). In contrast, the enzyme Tp-Man5 was not observed in the gel; however, some residual activity (0.6 U/ml) was linearly detected from 18 to 120 h, suggesting that this thermophilic enzyme is resistant to protease degradation (Figure 1C; *middle panel*). Notably, the enzymatic activity of this GH5 mannanase was measured at a high temperature (87 °C), avoiding the possibility of false-positive results. Protein secretion driven by strong promoters (such as *glaAp*) is often affected by extracellular proteolysis. This is a very common phenomenon, occurring in filamentous fungi that secrete native proteases, or in response to glucose starvation or increased pH (Budak et al., 2014; Segato et al., 2012; Yoon et al., 2011).

**Figure 1.**
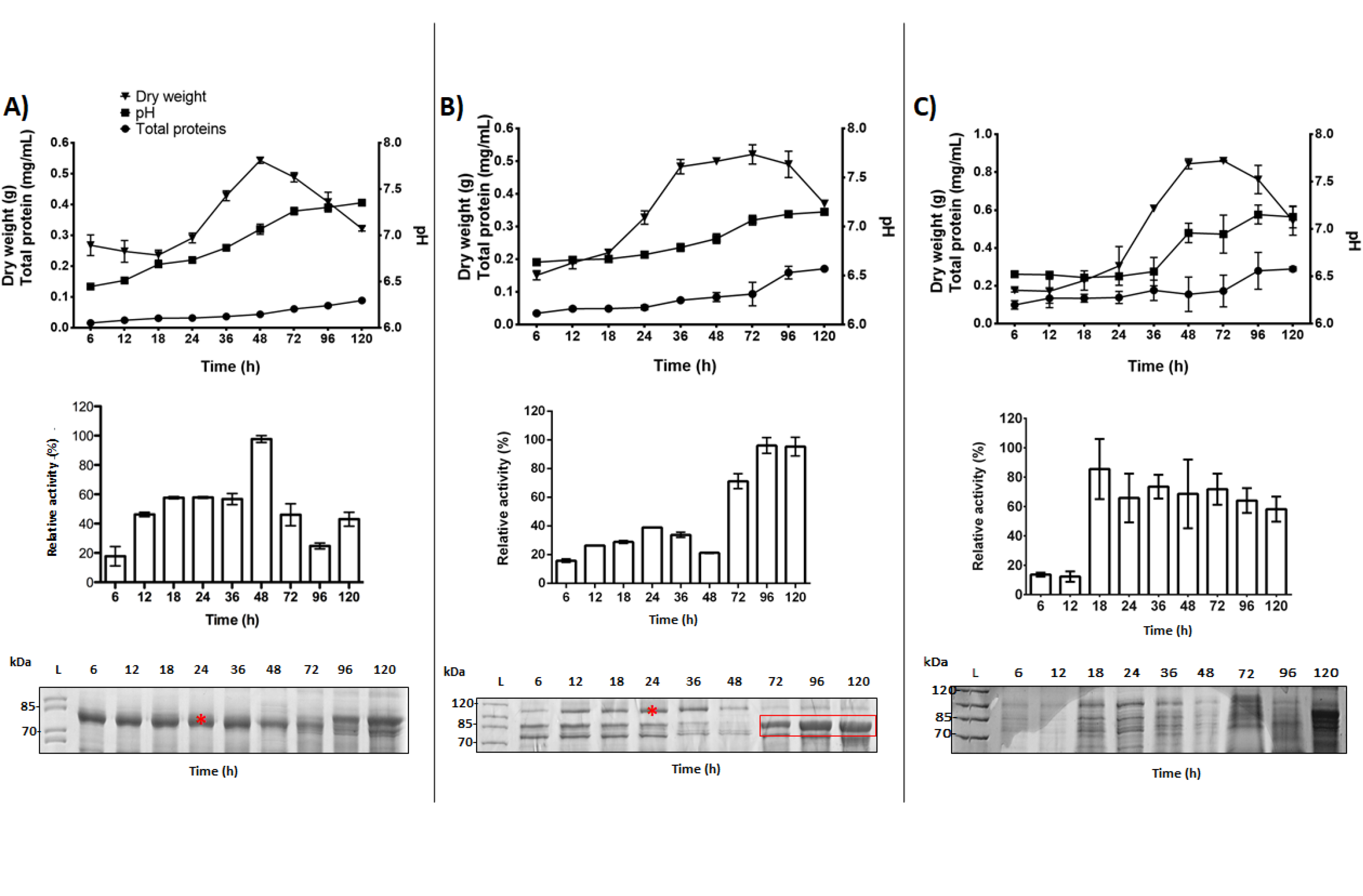
Physiological parameters and heterologous protein production by *A. nidulans* recombinant strains. A) *A. nidulans*_AbfA_, B) *A. nidulans*_BglC_, C) *A. nidulans*_Tp-Man5_. Physiological parameters (top): mycelium dry weight (g), final pH and extracellular proteins (mg/mL). Enzymatic activity of the *A. nidulans* strains during different cultivation periods (center). The highest enzymatic activity in each culture was considered to be 100%. SDS-PAGE of the heterologous proteins secreted by the recombinant *A. nidulans* strains during different cultivation periods (bottom). Ten micrograms of protein was applied in each lane. The target proteins are indicated by an asterisk. The red box indicates a native intracellular beta-glucosidase detected after 48 h of cultivation. The band corresponding to the Tp-Man5 protein was not detected by SDS-PAGE (bottom at right). Error bars indicate the standard deviation of three replicates.

To investigate whether the low secretion of the BglC and Tp-Man5 bands was caused by folding and/or secretion impairment, protein profiles and enzymatic activity assays were analyzed in the intracellular fraction (**Figure S1**). The overexpression of recombinant proteins may cause their intracellular accumulation, suggesting breakdown or overloading of the secretory pathway (Sims et al., 2005). However, BglC and Tp-Man5 were not detected in the intracellular fraction, indicating that these proteins did not accumulate inside the cell.

In this first analysis, we observed that recombinant protein secretion levels were higher for AbfA and BglC than for Tp-Man5, which showed residual activity in *A. nidulans*.

### Heterologous genes were highly expressed in *A. nidulans*

To investigate whether the differences in recombinant enzymes production were linked to mRNA abundance, the expression of *abfA, bglC* and *tp-man5* was measured by qPCR. The *abfA* and *bglC* genes were more highly expressed than the endogenous reference gene *(tubC)*, while the levels of *tp-man5* were similar to those of *tubC* (Figure 2A).

**Figure 2.**
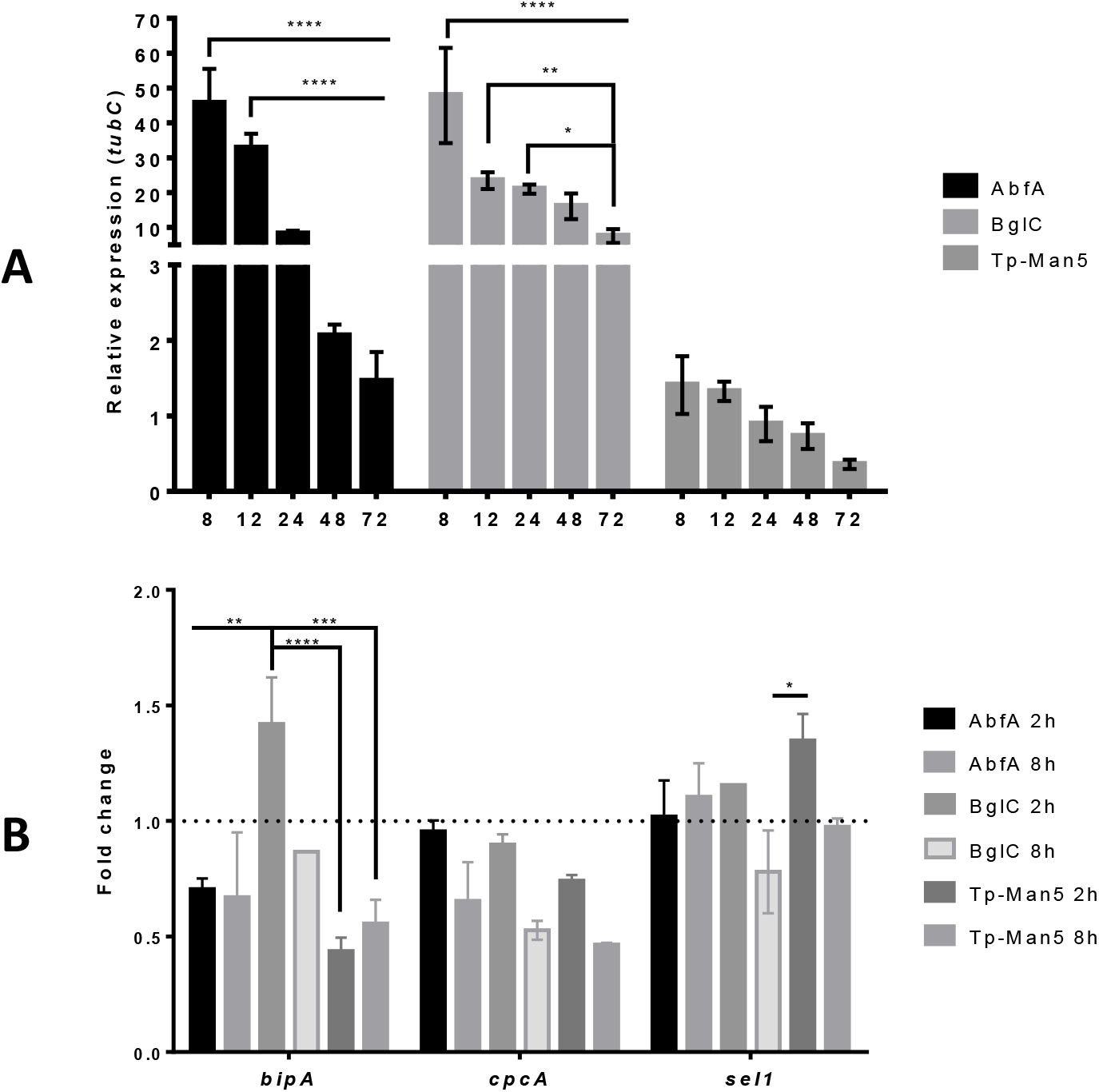
Gene expression profiles of *A. nidulans* recombinant strains. Expression levels were determined by qRT-PCR for different cultivation periods and expressed on a logarithmic scale as fold changes. A) Gene expression profile of the heterologous genes *abfA, bglC* and *tp-man5*. B) Expression of *bipA, cpcA* and *sel1* in *A. nidulans*_AbfA_, *A. nidulans_BglC_* and *A. nidulans*_Tp-Man5_. *tubC* was used as the reference gene, and the strain *A. nidulans*_A773_ was the experimental control. The ΔΔCt method was used to calculate gene expression. Asterisks indicate significantly different results (twoway ANOVA and Bonferroni posttest p-value<0.05). The strains were cultivated as described in the Materials and Methods section. The results are expressed as the mean values of three biological replicates and the error bars indicate standard deviation.

The production of recombinant proteins in filamentous fungi controlled by strong promoters activates some regulatory genes of protein quality control and UPR (Liu et al., 2014; Nevalainen and Peterson, 2014; Pakula et al., 2003). The UPR represents an adaptive response to restore cellular homeostasis, triggered by ER stress (Guillemette et al., 2011). The basic sensing pathway to detect ER stress or an increase in the folding load is highly conserved from yeast to humans. The accumulation of misfolded proteins stimulates Ire1 autophosphorylation and dimerization, triggering the unconventional splicing of an intron in the *hacA^u^* to create the transcriptionally active form *hacA^i^*(Guillemette et al., 2011). HacA is a conserved bZIP transcription factor in eukaryotic cells, regulating gene expression in response to various forms of secretion stress and as part of secretory cell differentiation (Carvalho et al., 2012).

UPR target genes such as *bipA, cpcA* and *sel1/ubx2* were quantified as a function of time (Figure 2B) (Cerqueira et al., 2014; Heimel, 2014; Sims et al., 2005; Wood et al., 2012). The putative ubiquitin-protein ligase-encoding gene *sel1/ubx2* was slightly overexpressed in the recombinants strains at 2 h and showed a tendency to normalize after 8 h. Likewise, *bipA* was overexpressed in the BglC-producing strain at 2 h and returned to basal levels at 8 h. *cpcA* was not overexpressed in any of the recombinant strains.

In *Trichoderma reesei*, the production of recombinant proteins moderately induced the overexpression of UPR genes only at early cultivation stages (from 1 to 12 h), and after 12 h, the expression of these genes returned to basal levels (Wang et al., 2014). Heterologous expression of the bacterial xylanase B (*xynB*) in *A. niger* also resulted in lower mRNA levels of some UPR target genes (*gla, bip1*, and *hac1*) (Zhang et al., 2008).

An indirect relationship between mRNA levels and the production of recombinant proteins was also reported for cellobiohydrolases (CBHs) from *Aspergillus terreus* and *T. reesei* expressed in *Aspergillus carbonarius* (Zoglowek et al., 2015). These authors speculated that proteolytic degradation could be one reason, possibly along with other factors such as incorrect folding, posttranslational processing, and impairment of intracellular transport (Zoglowek et al., 2015). This indirect relationship is frequently reported, suggesting that transcription is not a bottleneck in these systems.

In an attempt to identify the cellular processes altered during the adaptation of *A. nidulans* to the overexpression of recombinant genes, we analyzed the three recombinant strains by RNA-seq. We defined 2 and 8 h as standard time points for RNA-seq, based on the time course of recombinant protein production, the maltose consumption profile (**Figure S2**) and qPCR data.

### Global transcriptional response to heterologous protein production

Differential expression was determined by pairwise comparisons of the recombinant strains and the control strain *A. nidulans*_A773_. Four hundred seventy-six DE genes were identified by applying the pipeline and thresholds described in the Methods section (Figure 3). Thirty out of 476 DE genes were common to the three recombinant strains. *A. nidulansAbfA* and *A*. nidulansTp-Man5 also had 74 DE genes in common (Figure 3 **and Figure S3**).

To explore the cellular processes enriched by the production of recombinant proteins in *A. nidulans*, the annotation of DE genes was performed by The Functional Catalogue (FunCat) (Ruepp, 2004). The most significant functional category was “metabolism”, followed by “protein with binding function or cofactor”, “cellular transport” and “cell defense” (Figure 4). These four categories were enriched in all the recombinant strains and were also reported in other *Aspergillus* species overproducing recombinant proteins (Kwon et al., 2012; Liu et al., 2014; Zhou et al., 2016). The overexpression of glucoamylase (GlaA) in *A. niger* resulted in the enrichment of “translocation”, “protein glycosylation”, “vesicle transport” and “ion homeostasis” processes (Kwon et al., 2012). The functional categories “cellular transport”, “amino acid metabolism”, “aminoacyl -tRNA biosynthesis” and “metabolism” were overrepresented in an *Aspergillus oryzae* recombinant strain expressing a constitutively active form of *hacA*, indicating its importance for UPR, fungal growth, and physiology (Zhou et al., 2016).

**Figure 3.**
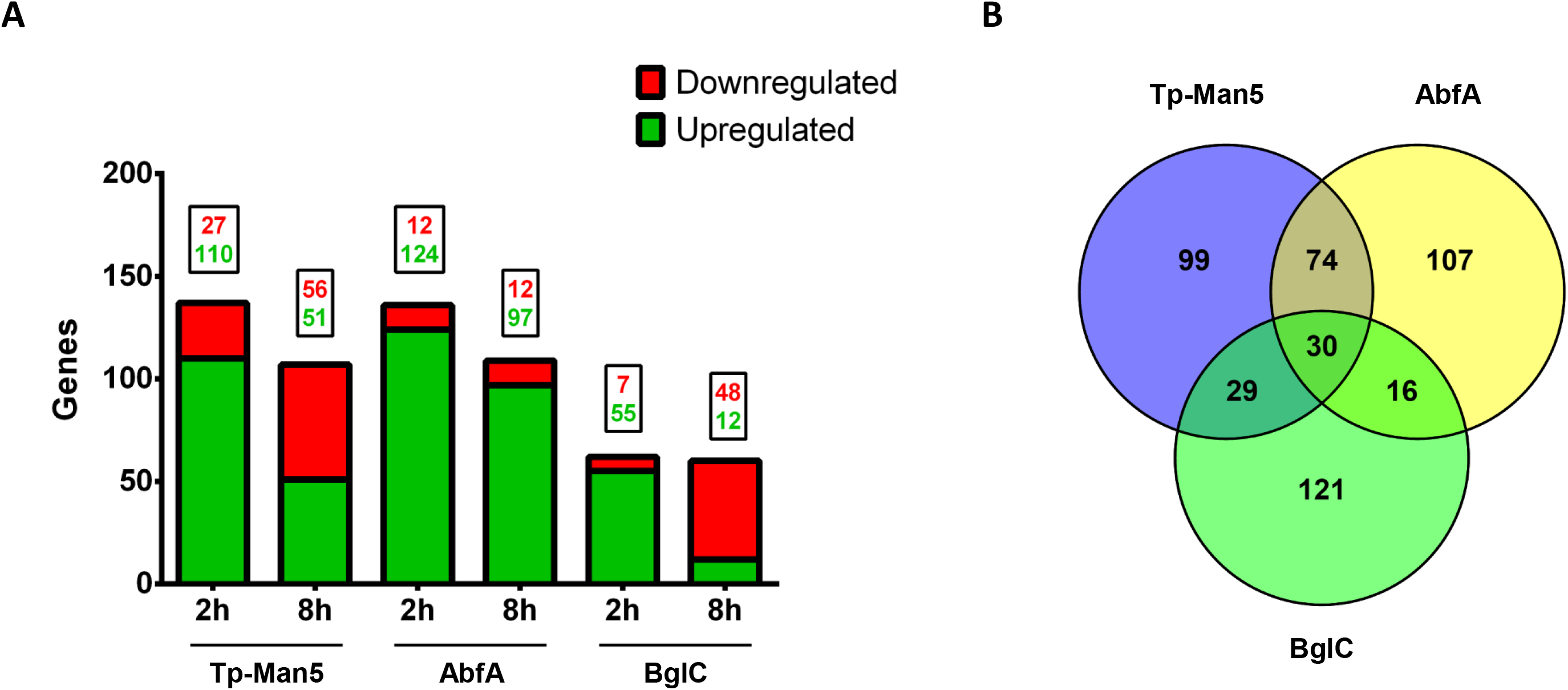
Global analysis of DE genes. A) Bar chart representing DE genes in the *A. nidulans*recombinant strains. Rectangles above the bars indicate the number of up- and downregulated genes. RNA-seq data for each recombinant strain were compared to that of the control strain *A. nidulans*_A773_, and a log 2 fold change ≤ −2.0 or ≥ 2.0 and adjusted p-value ≤ 0.01 were used as thresholds. B) Venn diagram representing DE genes analyzed by RNA-seq at 2 and 8 h of cultivation for the three *A. nidulans* recombinant strains. The Venn diagram was created using the online tool Venny 2.1.

**Figure 4.**
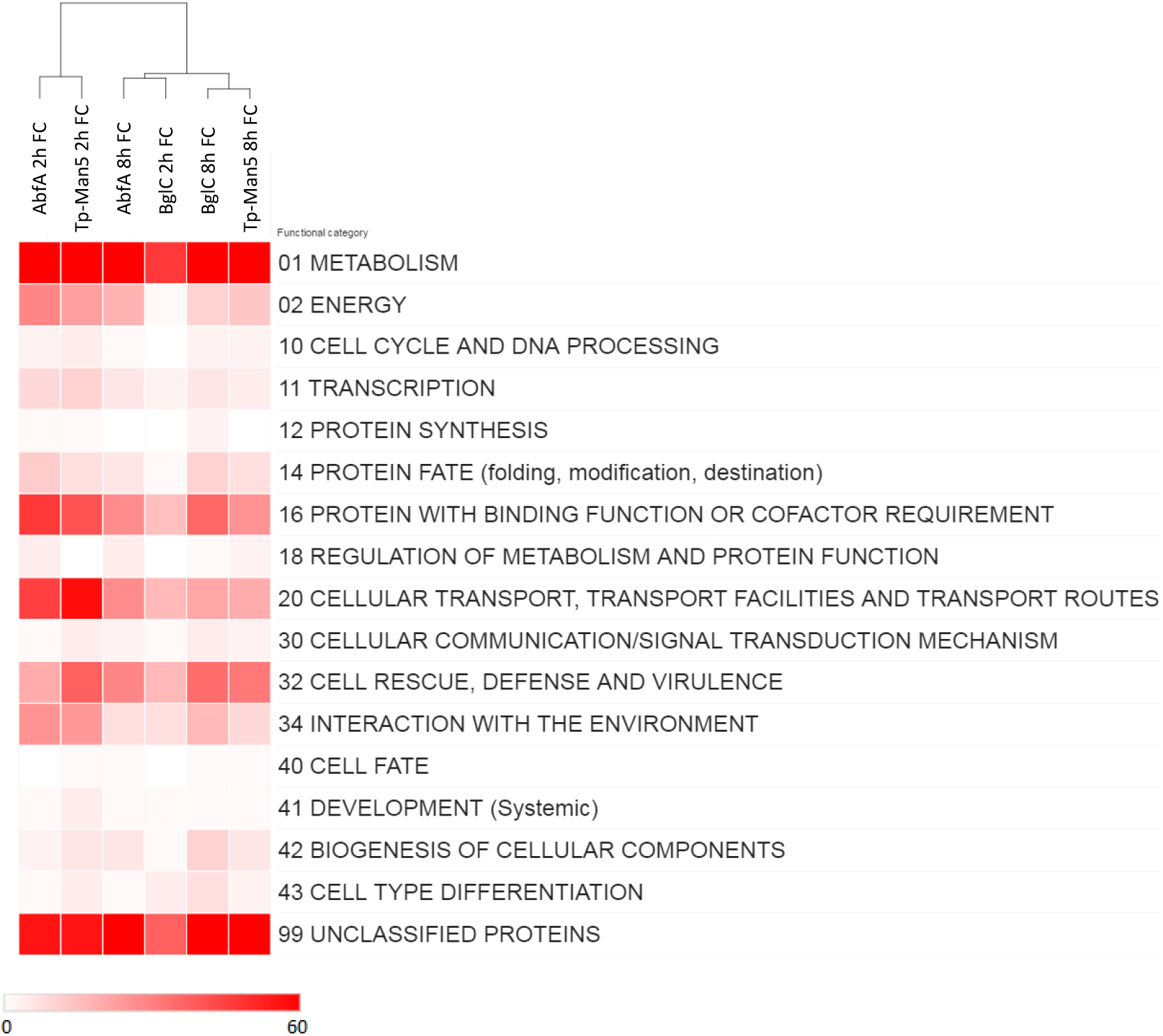
Functional annotation of the transcriptomic data for *A. nidulans* recombinant strains. MIPS FunCat term annotations of *A. nidulans*_Tp-Man5_, *A. nidulans*_AbfA_ and *A. nidulans*_BglC_ shows a broad variety of biological processes associated with DE genes of the *A. nidulans* recombinant strains. Analysis of the functional categories from global transcriptomic data was performed with FunCat. Functional categories in the heatmap represent the number of DE genes found in the transcriptomic data based on reference genome annotation. The heat map was created with online software from the Broad Institute, Morpheus.

Moreover, three patterns of expression were observed for the 30 DE genes in the three recombinant strains: genes upregulated at 2 and 8 h, genes downregulated at 2 h and upregulated at 8 h, and genes upregulated at 2 h and downregulated at 8 h (Figure 5, **Table S1 and S2**). These genes represent biological processes such as sexual sporulation, defense, detoxification and secondary metabolism, which are likely to constitute a common response of *A. nidulans* cells under recombinant protein production independent of the protein’s features, such as size, the complexity of folding and posttranslational modification pattern.

**Figure 5.**
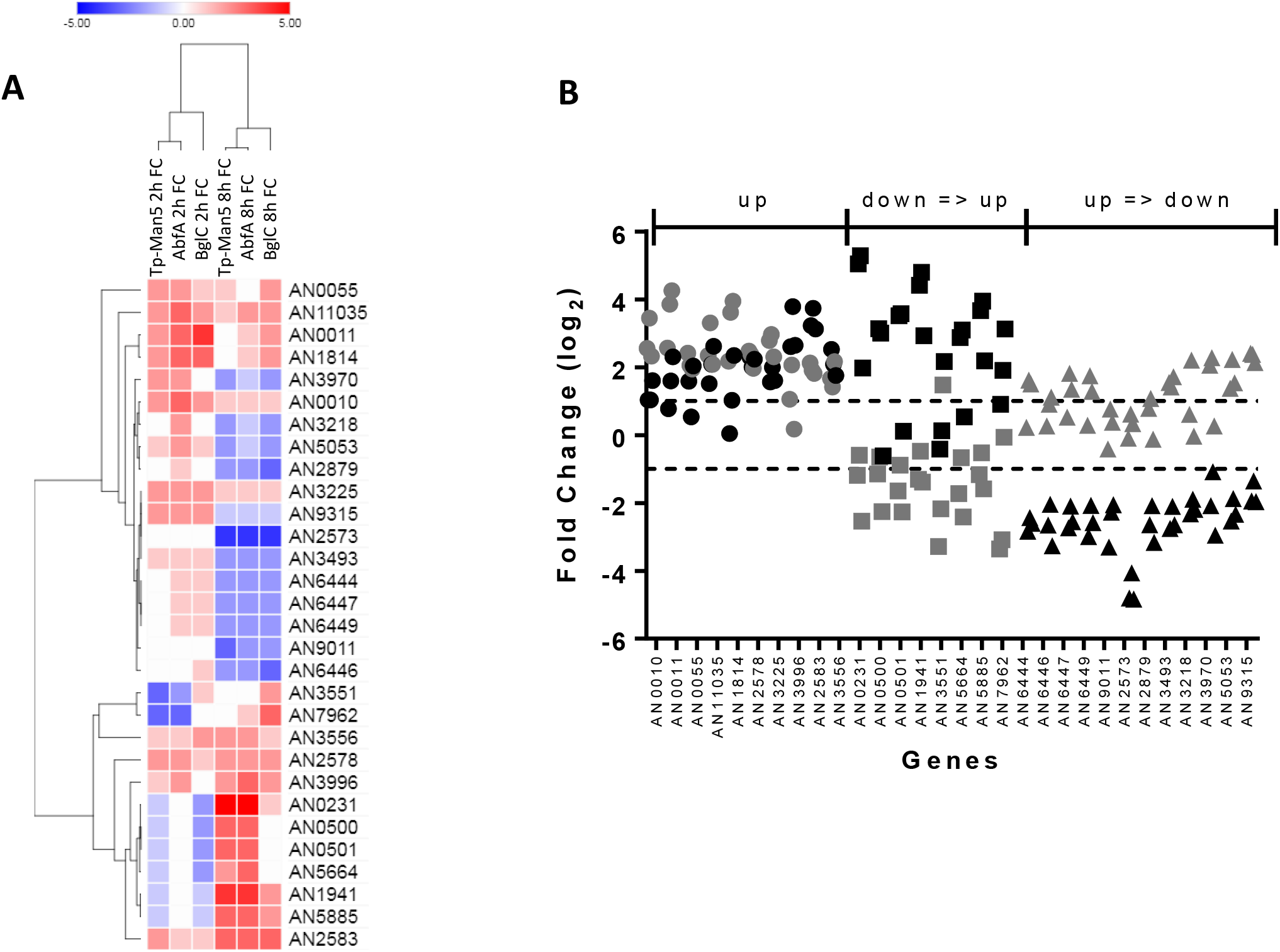
DE genes common to all recombinant strains. (A) Hierarchical clustering fell into three major clusters. (B) A schematic representation of each pattern of transcriptional change: up (increased at 2 and 8 h), down to up (decreased at 2 h and increased at 8 h), and up to down (increased at 2 h and decreased at 8 h). Circles: 10 genes; squares: 8 genes; triangles: 12 genes. Gray symbols: 2 h. Black symbols: 8 h.

To evaluate the canonical pathway of UPR, we analyzed the ratio of *hacA* splicing in *A. nidulans* recombinant strains based on the RNA-seq data (**Figure S4**). The levels of *hacA* splicing were higher for AbfA and Tp-Man5 than for BglC at 2 h, suggesting the presence of misfolded proteins. After 8 h, the levels of *hacA* splicing for all the recombinant strains were lower than those for the control strain, showing normalization of the UPR canonical pathway. This result suggests either no accumulation of the BglC misfolded form or the presence of an alternative ER stress and UPR pathway. The RNA-seq and qPCR data showed an acceptable correlation (**Figure S5**). Wang et al (2014) analyzed *hac1* levels in the *T. reesei* Rut C30 and QM9414 strains. The transcript levels of *hac1* increased earlier in Rut C30 (1 h of induction), corroborating our data (Wang et al., 2014).

Based on the RNA-seq data for the recombinant strains, we observed a globally upregulated transcriptional response. Moreover, biological processes related to metabolism, protein with binding function and cellular transport were enriched. Unconventional splicing of *hacA* was observed in *A. nidulans_AbfA_* and *A. nidulans_Tp-Man5_*, indicating some level of ER stress. The global analysis showed mild stress at 2 h after the induction of heterologous protein production, which was normalized after 8 h.

### Differential expression of genes related to the secretion pathway

The production of heterologous proteins is not only impaired by the low expression of heterologous genes but can also be reduced by problems with secretory pathway posttranslational processing (Yoon et al., 2010). Many studies have attempted to understand the high capacity of *Aspergillus* for protein production, primarily at the transcriptional level (Carvalho et al., 2012; Guillemette et al., 2007; Kwon et al., 2012; Liu et al., 2014; Sims et al., 2005). These studies identified important genes in different stages of the protein production pathway, such as translocation, folding, cargo transport and exocytosis (Schalén et al., 2016). Liu et al. (2014) listed a set of genes involved in the secretion pathway of *A. oryzae* using the secretory model *S. cerevisiae* as a scaffold. Based on this list, we defined the homologous or best-hit genes in *A. nidulans* by using the AspGD data, grouping a set of 374 genes (**Table S3**).

Seventeen genes were DE in the recombinant strains (**Table S4**; Figures 6A and B). The most highly represented categories were “stress response”, “protein folding and stabilization” and “unfolded protein response”, with 10, 8 and 8 genes, respectively (Figure 6C). To gain insights into genetic interactions, ten different RNA-seq networks were calculated to show DE genes, and each of the 17 genes involved in protein secretion was present in no more than one network (**Figure S6**). The number of nodes and edges of the networks varied from 40 and 507 to 1155 and 276232, respectively. Biological process enrichment was carried out for entire networks as well as specific genes of interest and their close neighbors.

**Figure 6.**
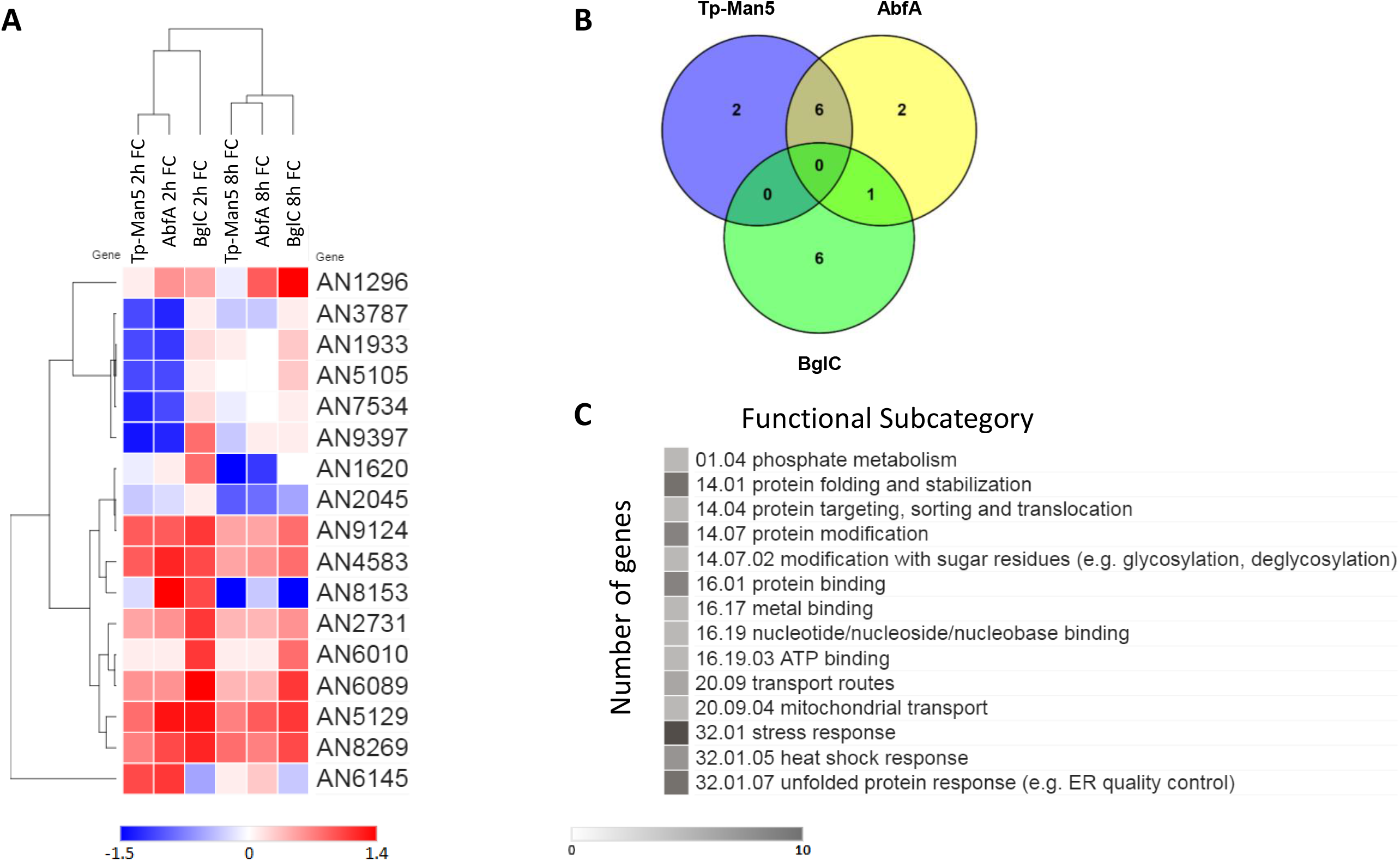
Analysis of genes with predicted functions in the secretory pathway. A) Heatmap representing the 17 DE genes with predicted functions in the in the secretion pathway in *A. nidulans*.B) Venn Diagram. No gene was common to all recombinant strains. The Venn diagram was created using the online tool Venny 2.1. C) Functional annotations of DE genes. The analysis of functional categories was performed with FunCat. The values represent the number of altered genes found (abs set) in the transcriptomic data based on the values of the reference genome annotation. All functional subcategories with at least five DE genes are represented. *A. nidulans* strains were cultivated as described in the Materials and Methods section. Heatmaps were created with online software from Broad Institute, Morpheus.

**Figure 7.**
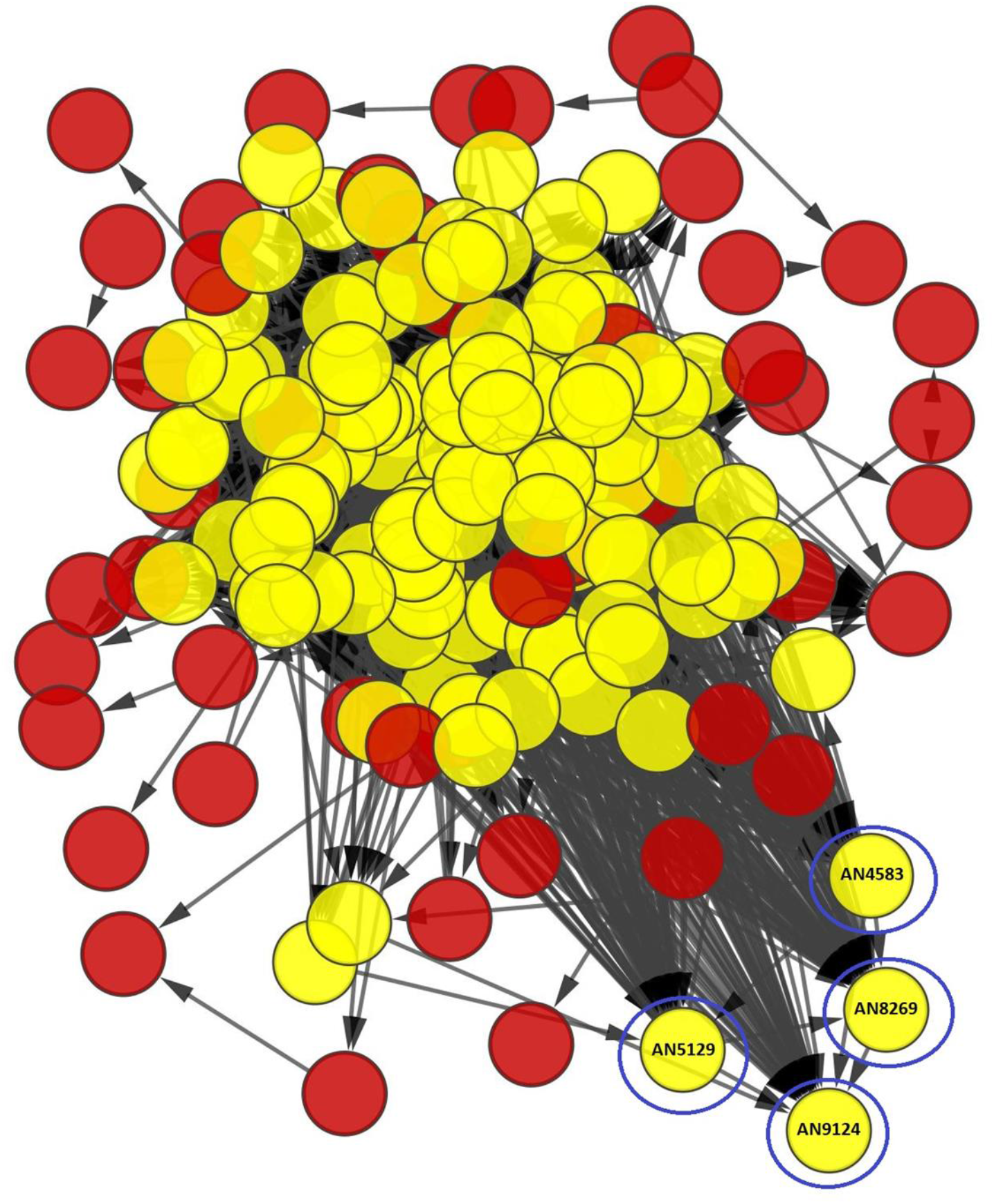
RNA-seq network. DE genes in which “protein folding” was observed as an enriched biological process. Yellow circles represent DE genes (AN9124, AN4583, AN5129 and AN8269) with predicted functions in the secretion pathway and their first interactions. Genes of interest are found within blue circles.

Among the 17 genes mentioned previously, AN1296, AN3787, AN1620, AN2045, AN8153, AN6010 and AN6145 were found in different networks. Regarding biological process enrichment for all networks, a *p*-value ≤ 10^-3^ was considered significant. In the networks containing AN1296 and AN6010, no statistically significant enriched biological processes were found, but for the other five genes, different processes were enriched, including amino acid, carboxylic acid, aldehyde and aromatic amino acid catabolic processes; chromosome organization; cellular response to DNA damage stimulus; cellular response to stress; filamentous growth; hyphal growth; cell development; biosynthetic process; metabolic process; precursor metabolite energy; and proton transport.

Although these seven aforementioned networks contained only one gene of interest, three networks included more than one gene. AN2731 and AN6089 were found in the same network and were upregulated in *A. nidulans*_BglC_. Both genes are predicted to encode different heat shock proteins. The genes AN1933, AN5105, AN7534 and AN9397 were also found in the same network. Interestingly these four genes were downregulated in *A. nidulans*_Tp-Man5_ and *A. nidulans*_AbfA_ (2h). However, no biological processes were found to be enriched. In addition, the deletion of orthologs of AN1933, AN5105 and AN9397 was not lethal in *S. cerevisiae*, although growth defects or decreased levels of protein were observed in specific situations (Han et al., 2010; Kriangkripipat and Momany, 2009; Valkonen et al., 2003).

Finally, AN9124, AN4583, AN5129 and AN8269 were DE and grouped within the same network (Figure 7). AN4583 encodes a putative peptidyl-prolyl cis-trans isomerase D, and the transcriptional level of its ortholog (CPR7) in *S. cerevisiae* was increased during UPR. This enzyme mediates signaling or conformational changes in the chaperone Hsp90, and its deletion resulted in growth defects (Zuehlke and Johnson, 2012). The other genes (AN9124, AN5129 and AN8269) encode heat shock proteins involved in protein folding during stress. The deletion of their orthologs in *S. cerevisiae* was not lethal (Chang et al., 1997; Floer et al., 2008; Matsumoto et al., 2005). Interestingly, this was the only network in which “protein folding” was observed as an enriched biological process, for the whole network as well as for the specific genes and their close neighbors. Thus, these four genes are potential targets for genetic manipulation to improve heterologous protein production in *A. nidulans*.

The information generated from the transcriptomic analysis is highly important for selecting the best targets for genetic engineering of the secretory pathway of *Aspergillus* strains. The deletion or overexpression of such genes can interfere with enzyme production. For instance, overexpression of the Rab GTPase *rabD* in *A. nidulans*, which is involved in vesicle-mediated exocytic secretion and autophagy, improved the production of a recombinant glucoamylase by 40%. On the other hand, the overexpression of other genes, such as AN6307 and AN4759, which are involved in the transport of proteins from the ER to the Golgi and from the Golgi to the plasmatic membrane, considerably reduced recombinant enzyme secretion (Schalén et al., 2016).

The deletion of AN6305 (PkaA), which encodes cAMP-dependent protein kinase A, enhanced the secretion of hydrolytic enzymes in *A. nidulans*. The knockout strain showed increased hyphal branching, higher expression of SynA, involvement in secretion, and increased expression of genes associated with mitochondrial function, fatty acid metabolism, and the use of cell storage (de Assis et al., 2015). In a different work, the deletion of an α-amylase, which is the most highly secreted protein in *A. oryzae*, alleviated UPR induction and resulted in the overexpression of a mutant 1,2-α-D-mannosidase (Yokota et al., 2017). In addition, the double deletion of CreA and CreB, which are involved in the regulatory machinery of carbon catabolite repression, in *A. oryzae* resulted in more than 100-fold higher xylanase and β-glucosidase activities than those in the wild-type strain (Ichinose et al 2017).

## CONCLUSIONS

The production of recombinant proteins in filamentous fungi remains a “black box” because biotechnology companies have carried out part of the scientific development in this field. All insights at the genomic, transcriptomic and proteomic level are welcome and may reveal a pipeline for genetic manipulation to generate “superior” strains for enzyme production.

In our manuscript, the first important result was the analysis of a thermophilic gene in *A. nidulans*. Even a gene from a phylogenetically distant bacterium was properly expressed in *A. nidulans*, despite low levels of protein production. Moreover, for BglC and Tp-Man5, the low levels of production are probably related to translation instead of secretion since these proteins were not found to be trapped inside the cell. Therefore, the first part our results showed the indirect relationship between mRNA expression and protein production, highlighting a thermophilic gene. Another interesting result was that, despite the distinct features of the recombinant proteins, 30 DE genes were common to all the recombinant strains. A likely conclusion is that these genes represent a general response to the expression of heterologous genes. Even though we did not design knockout mutants to address the functions of these 30 genes in protein production/secretion, the identification of these genes themselves offers new targets for genetic manipulation and strain improvement.

We also showed early activation of the canonical UPR pathway by *hacA* alternative splicing that was normalized after 8 h, although not in the strain expressing BglC, suggesting either that there was no accumulation of the BglC misfolded form or the presence of an alternative ER stress and UPR pathway. Finally, in order to focus our analysis on the secretion pathway, a set of 374 genes was further evaluated. Seventeen genes were common to all the recombinant strains, suggesting again that these genes represent a general response of *A. nidulans* cells to the overexpression of recombinant genes, even a thermophilic gene. Additionally, we reported the possible genetic interactions of these 17 genes based on coexpression network calculations.

Our data highlight the complexity of producing heterologous enzymes in *A. nidulans*. The response to heterologous protein production is nongeneric, and improvements in the production of one target protein are not necessarily transferable to other strains. Thus, this study may provide a genetic and cellular background and targets for genetic manipulation to improve protein secretion by *A. nidulans*.

## MATERIALS AND METHODS

### Strain maintenance and cultivation conditions

*A. nidulansA773* (pyrG89;wA3;pyroA4) was purchased from the Fungal Genetics Stock Center (FGSC, St Louis, MO). The strains *A. nidulansAbfA, A. nidulansBglC* and *A*. nidulansTp-Man5 were previously transformed and tested by our research group, as described by Segato 2012 (Segato et al., 2012). We purchased 5-fluorotic acid (5-FOA) from Oakwood Products Inc (NC9639762), and all other chemicals were from Sigma–Aldrich, Megazyme and Fisher Scientific.

Vegetative cultures and conidia production cultures were prepared by cultivation in minimal medium as described by Clutterbuck (Clutterbuck, 1992) and Pontecorvo (Pontecorvo et al., 1953). *A. nidulans* strains were cultivated in minimal medium containing 5% 20× Clutterbuck salts, 0.1% 1000× vitamins, 0.1% 1000× trace elements, pH 6.5 and supplemented with pyridoxine (1 mg/L), uracil/uridine (2.5 mg/L each) whenever required. Minimal medium was supplemented with 1% glucose for mycelium growth (growth medium) or 2% maltose and 250 mM HEPES buffer (Sigma–Aldrich) for promoter activation and protein production (induction medium). Incubation was carried out at 37 °C.

### Production and secretion of client proteins

Pre-cultures were prepared by inoculating fresh 10^8^ spores/ml in 30 mL of minimal medium supplemented with 1% (w/v) glucose for the mycelium growth for 48 h at 150 rpm and 37 °C. Strains cultivation were performed in biological triplicates. The mycelium was washed with autoclaved distilled water, filtered with Miracloth, and then transferred to the induction medium. After defined periods for protein induction, the mycelium was collected, Miracloth filtered, dried, frozen in liquid nitrogen and stored at −80 °C. The medium was collected, centrifuged at 10,000 × g for 10 min prior to concentration by ultrafiltration (10 kDa cutoff Amicon), quantified by the Bradford method(Bradford, 1976), purity was assessed by SDS-PAGE (Shapiro and Maizel, 1969) and then used for biochemical studies.

### Analytical assays

The enzymatic activity of beta-glucosidase GH3 (BglC) was determined with 4-nitrophenyl-β-D-glucopyranoside (Sigma–Aldrich) and that of alpha-arabinofuranosidase GH51 (AbfA) with 4-nitrophenyl-α-L-arabinofuranoside (Sigma–Aldrich). The enzymatic activity assays were performed by adding 40 μL of extracellular crude extract to 50 μL of each substrate at 5 mM in 10 μL of 100 mM ammonium acetate buffer (pH 5.5) followed by incubation for 45 min at 50 °C. The enzymatic reactions were stopped with sodium carbonate. The absorbance was read at 405 nm with a Multimode Infinite M200 Reader (Tecan, SC), and activity was calculated using p-nitrophenol as the standard.

The enzymatic activity of thermophilic mannanase GH5 (Tp-Man5) was determined with locust bean gum (Sigma–Aldrich) as the substrate. The reaction was performed by adding 1 μg of proteins from concentrated extracellular crude extract to 50 μL of 1% (w/v) substrate in 50 mM ammonium acetate buffer (pH 5.5) and incubating for 10 h at 87 °C. The release of reducing sugars was measured with DNS acid (Miller, 1959), and absorbance was read at 540 nm with a Multimode Infinite M200 Reader (Tecan, SC) and compared to a glucose standard curve. Statistical calculations and plotting were carried out with GraphPad Prism v 5.0 Software (California, US).

The concentration of maltose was detected by high-performance liquid chromatography (HPLC) Agilent Infinity 1260 with IR detector 50C, Aminex column HPX-87P 300 mm × 7.8 mm at 50 °C and 0.5 mL/min of ultrapure Milli-Q water as eluent phase.

### Intracellular protein analysis

Intracellular proteins were extracted from frozen mycelium after grounding in liquid nitrogen to a fine powder using a mortar and a pestle. The powder was gently suspended in 5 mL of extraction buffer (20 mM Tris-HCl pH 8.0; 0.05 *%* (w/v) Triton™ X-100 (Sigma–Aldrich), and 150 mM NaCl, containing protease inhibitors (2 mM PMSF and Protease Inhibitor Cocktail N221-1mL from Amresco), and centrifuged twice at 8000 rpm for 15 min at 4 °C; then, the supernatant was collected for SDS-PAGE analysis (Shapiro and Maizel, 1969).

### RNA extraction

Total RNA was extracted from frozen mycelium using the Quick-RNA™ MiniPrep kit (Zymo Research) according to the manufacturer’s instructions. The integrity of extracted RNA was evaluated (RIN ≥ 8) on the Agilent Bioanalyzer 2100 and quantified using an ND-1000 NanoDrop (Thermo Scientific) spectrophotometer. Total purified RNA was stored at −80 °C until further processing.

### RNA-seq data analysis

For RNA-seq sample preparation, total RNA was obtained from the four *A. nidulans* strains (three recombinant strains and the parental strain) cultivated for 2 and 8 h in biological triplicate, resulting in twenty-four samples for library preparation using the TruSeq Stranded mRNA Library Prep Kit v2 (Illumina) according to the manufacturer’s instructions. Sequencing libraries were prepared and sequenced using Illumina HiSeq 2500 at the CTBE NGS sequencing facility. Approximately 148 million 100-bp paired-end reads were obtained for the *A. nidulans*_A773_, *A. nidulans*AbfA, and *A. nidulans*_BglC_ strains and 130 million reads were obtained for the *A. nidulans*_Tp-Man5_ strain, representing a total of 115 GB. Reads were quality-checked and filtered using FASTQC and Trimmomatic (Bolger et al., 2014), respectively, and rRNA contamination was assessed and removed using sortmeRNA (Kopylova et al., 2012). The rate of rRNA contamination was lower than 16% in all samples, except in one 8 h AbfA replicate (**Table S5**). QC reads were aligned to the *Aspergillus nidulans* genome available at AspGD (http://www.aspgd.org/) using TopHat2 (Kim et al., 2013), and the concordant pair alignment rate varied between 84 and 93%. An acceptable replicate agreement was obtained for all conditions, as shown by PCA analysis and clustering of the samples data.

For differential expression analysis, pairwise comparisons between each strain and the control strain *A. nidulans*_A773_ were performed using the DESeq2 R/Bioconductor package (Love et al., 2014), applying a log2 fold change ≥ 2 or ≤ −2 and an adjusted p-value ≤ 0.01 as thresholds. For secretion pathway analysis, a log2 fold change ≥ 1 or ≤ −1 and an adjusted p-value ≤ 0.01 were used as thresholds. Graphs were constructed using GraphPad Prism v 5.0 Software and an online tool for Venn diagrams, Venny 2.1 (http://bioinfogp.cnb.csic.es/tools/venny/).

### Functional annotation

For functional annotation of DE genes identified by RNA-seq analysis, lists of genes were generated by filtering the results with log 2 FC ≤ −1.0 or ≥ 1.0 and adjusted p-value ≤ 0.01. The gene expression values of the recombinant strains were compared to the control strain *A. nidulans*_A773_. The gene lists obtained were then used for functional annotation using FunCat (Functional Catalogue) (Ruepp, 2004).

The selected genes for annotation with FunCat were faced with the database of the reference genome of *A. nidulans* and applied statistical Fisher test with p-value ≤ 0.05 as a threshold. The enriched functional categories and subcategories were used for comparisons and graphics generation. Data analysis was performed using GraphPad Prism v 5.0 Software (California, US).

### Secretion pathway analysis

A list of 374 secretory pathway genes was generated based on Liu et al. (2014), who defined a list with the functional protein secretory components from *A. oryzae* using the secretory model *S. cerevisiae* as a scaffold (Liu et al., 2014). This list was further adapted to filamentous fungi by adding *A. oryzae* orthologs of the secretory components reported for other *Aspergillus* species, such as *A. nidulans* and *A. niger*. The homologous genes in *A. nidulans* reported in AspGD resources were then used to generate the list with 374 genes.

DE genes of the secretory pathway were functionally annotated with FunCat, and the most representative functional categories were used to build a secretory pathway profile of the three recombinant strains. To generate this profile, the FC values of the DE genes involved with secretion pathway were used to create heat maps using R.

### Quantitative real-time PCR (qRT-PCR) analysis

To quantify the expression of selected genes, qRT-PCR was performed. For this analysis, 2 μg of pure RNA was used for cDNA synthesis using the First Strand cDNA Maxima™ Synthesis Kit (Thermo Scientific), according to the manufacturer’s instructions. The cDNA was diluted 1/50 and used for analysis in the ViiA™ 7 Real-Time PCR System, using Maxima SYBR™ Green (Thermo Scientific) for signal detection, in accordance with the manufacturer’s instructions. The gene encoding β-tubulin (AN6838) was used as an endogenous control because it is a stable gene in filamentous fungi (Llanos et al., 2015).

The PCR conditions were as follows: 95 °C for 30 s, followed by 40 cycles at 95 °C for 10 s and 60 °C for 60 s. The melting curve was analyzed with ViiA™ 7 Software (Thermo Sci entific) to confirm the presence of only one amplicon, according to the Tm expected for each gene. Gene expression values were calculated according to the 2-ΔΔCT method (Livak and Schmittgen, 2001). Primers used in the qPCR experiments are described in **Table S6**. Data analysis was performed using GraphPad Prism v 5.0 Software (California, US).

### Weighted gene co-expression network analysis

We applied the WGCNA in order to identify groups (modules) of genes that showed highly co-expressed gene expression across the two treatments (2h and 8h) under a specific condition (Tp-Man5, BgCl and AbfA). The co-expression analysis was implemented with the WGCNA package in R (Langfelder and Horvath, 2008). Consensus WGCNA analysis consisted of construction of correlation matrices, which were then converted into adjacency matrices that retain information about the sign of the correlation (i.e., signed networks use a transformation of 0.5 ×(r+ 1)). The soft power threshold chosen was based on a measure of R2 scale-free topology model fit that maximized and plateaued well above 0.7 (i.e., soft power = 20 for both datasets). Modules were merged at a cut height of 0.2, and the minimum module size was set to 30. As a result, for both treatments, we got sets of genes (modules) that were highly co-expressed within the modules, but not necessarily between the modules. We identified 29 distinct co-expression modules that contain at least one secretion gene or a transcription factor.

### Functional enrichment analysis

To assess functional enrichment, Gene Ontology (GO) Biological Processes term and Kyoto Encyclopedia of Genes and Genomes (KEGG) pathway analyses of WGCNA network modules were performed using Cytoscape plug-in BinGO (Maere et al., 2005). These analyses provided a comprehensive set of functional annotation tools for investigators to understand the biological meaning behind large lists of genes.

## Data availability

The datasets generated and/or analyzed during the current study are available in the Gene Expression Omnibus (GEO) repository, GSE101522, https://www.ncbi.nlm.nih.gov/geo/query/acc.cgi?acc=GSE101522.

## ACKNOWLEDGMENTS

This research was supported in part by FAPESP (grant 2012/20549-4 to ARLD; 2014/06923-6 to FMS). We are grateful to the National Council for Scientific and Technological Development (CNPq) for the financial support (441912/2014-1 and 304816/2017-5 to ARLD). CRFT, MVR, MPZ and FC received FAPESP fellowships (2017/10083-1; 2013/24988-5; 2014/15403-6; 2014/23051-2). We thank the LNBio Mass Spectrometry staff for the assistance with LC-MS/MS and the CTBE High Throughput Sequencing and Robotics Laboratory (NGS) for the assistance with RNAseq.

## AUTHOR’S CONTRIBUTIONS

ARLD conceived and designed the experiments. FC, MVR, MPZ participated in the design of the study and performed the RNAseq experiments. GFP filtered the RNAseq raw data and generated the differential expressed genes lists. FC, MVR, MPZ, FJC, GFP and ARLD analyzed the RNAseq data. FC, FMS, FJC and CRFT helped to interpret the experimental data. FC and ARLD drafted the manuscript. ARLD, CRFT, FMS and GFP revised the manuscript. All authors read and approved the final manuscript.

